# SPECTRE: standardized global spatial data on terrestrial SPECies ThREats

**DOI:** 10.1101/2023.11.26.568516

**Authors:** Vasco V. Branco, Luís Correia, Pedro Cardoso

## Abstract

**Motivation:** SPECTRE is an open-source database containing standardised spatial data on global environmental and anthropogenic variables that are potential threats to terrestrial species and ecosystems. Its goal is to allow users to swiftly access spatial data on multiple threats at a resolution of 30 arc-seconds for all terrestrial areas. Following the standard set by Worldclim, this data allows full comparability and ease of use under common statistical frameworks for global change studies, species distribution modelling, threat assessments, quantification of ecosystem services and disturbance, among multiple other uses. A web user interface, a persistent online repository, and an accompanying R package with functions for downloading and manipulating data are provided.

**Main types of variable contained:** SPECTRE is a GIS product with 24 geoTiff raster layers (with plans to expand in the near future) with an approximate 1 km^2^ resolution.

## 1 BACKGROUND

Biodiversity and ecosystem services loss are ongoing issues that have taken the spotlight in several international agendas, such as the recently approved Kunming-Montreal Global Biodiversity Framework (CBD, 2023). Across all agendas the need for direct conservation actions and policy according to the best available knowledge is recognized. However, the status of biodiversity and the services it provides to humanity remain unquantified for the most part. This is at least partly due to the lack of data, but often the culprit is the low accessibility and comparability of the many data sources already available.

Standardised data on several variables of interest are already available or being mobilised through different platforms. These include species distributions through GBIF (Global Biodiversity Information Facility, 2022), and spatial or temporal trends of communities or populations through PREDICTS (Natural History Museum, 2022; Hudson *et al*., 2016) and BioTIME (Dornelas *et al*., 2018) respectively. Variables representing potential threats to species are also available from various sources, such as forest loss from ForestWatch (Global Forest Watch, 2014), or human density and built area from the Global Human Settlement Layer (Corbane, Florczyk, Pesaresi, Politis, & Syrris, 2018; Schiavina, Freire, & MacManus, 2019). Despite the availability of this data, these sources are not directly comparable and users that intend to use multiple threats in their analyses will most often have to go through the task of standardising data across a myriad of formats, resolutions and extents.

Here we present SPECTRE, a global spatial database of threats to terrestrial species and ecosystems. Our goals are fourfold: 1) compiling global spatial layers of several types of threats; 2) standardising them to a common extent and resolution, identical to existing and widely used climate layers; 3) make them easily and openly available to researchers and decision makers involved in species and ecosystem conservation; and 4) provide an online portal together with the tools to easily download and manipulate different layers in the R environment.

With these goals in mind, we have chosen WorldClim (Fick & Hijmans, 2017) as our template for SPECTRE. WorldClim is a longstanding, high quality dataset of global climate layers and has been used extensively in several fields, including species distribution modelling (SDM), understanding the effects of climate change on species, and more, with over 1000 citations per year. As such, by adopting WorldClim as our template, we guarantee the easy integration of SPECTRE in a myriad of studies involving both climate and biodiversity threats and straightforward comparison to the outcome of many studies.

It is our intention to make SPECTRE a continuously updated effort, advancing as needed through community suggestions and best available scientific knowledge on threats to species and ecosystems worldwide.

## 2 METHODS

### 2.1 Description of data

The first version of SPECTRE is composed of a collection of 24 standardised raster layers, each representing a potential threat to species and ecosystems (Fig. 1). The layers are classified (Table 1) first according to the list of direct pressures on biodiversity and sustainable use as proposed in the Convention on Biological Diversity (CBD, 2020a; CBD, 2020b) in their Strategic Goal B. Each is then further subdivided in sub-sections according to the International Union for the Conservation of Nature (IUCN) threat categories (IUCN, 2023).

**FIGURE 1.**
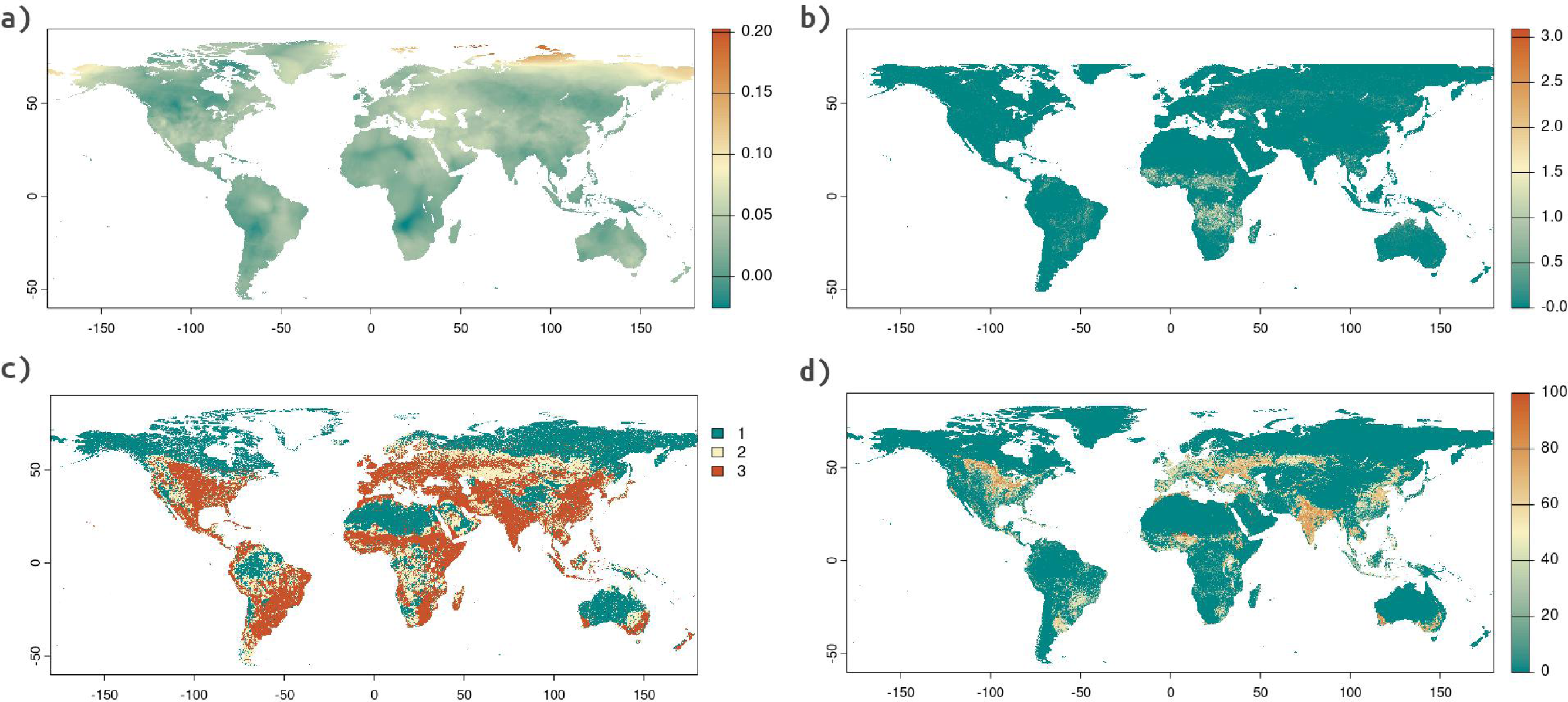
Example global raster layers for a) temperature trends (*5_1_TEMP_TRENDS)*, b) fire occurrence (*1_10_FIRE_OCCUR*), c) impact area (*1_7_IMPACT_AREA*) and d) crop percentage (*2_2_CROP_PERC_IIASA*). Layers in SPECTRE represent common threats to species, both directly as the destruction of species habitats, as well as indirectly, such as projected surface temperature changes for a given area. Note that some areas, such as polar regions, are not represented in all layers, depending on the original source.

**TABLE 1.**
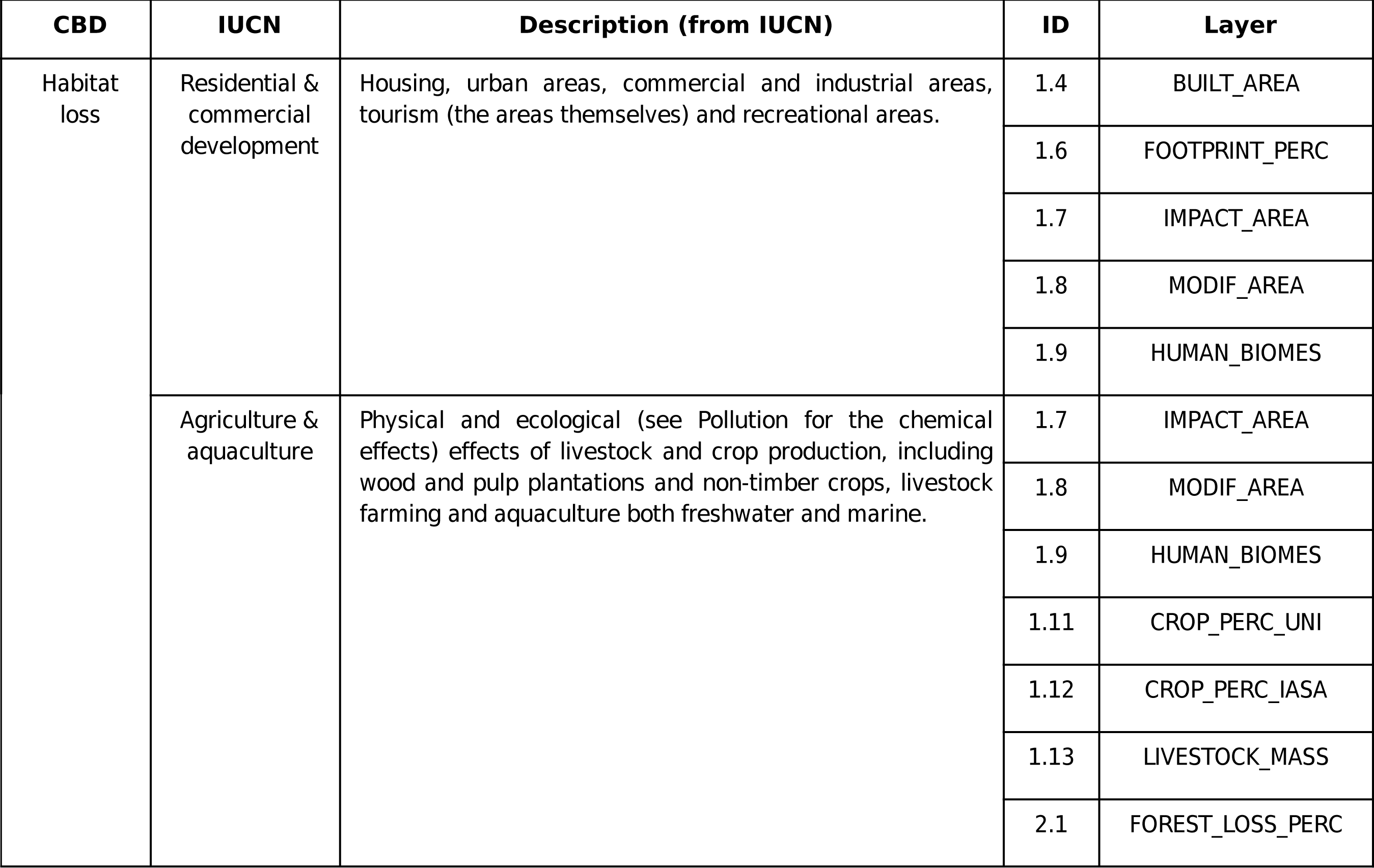
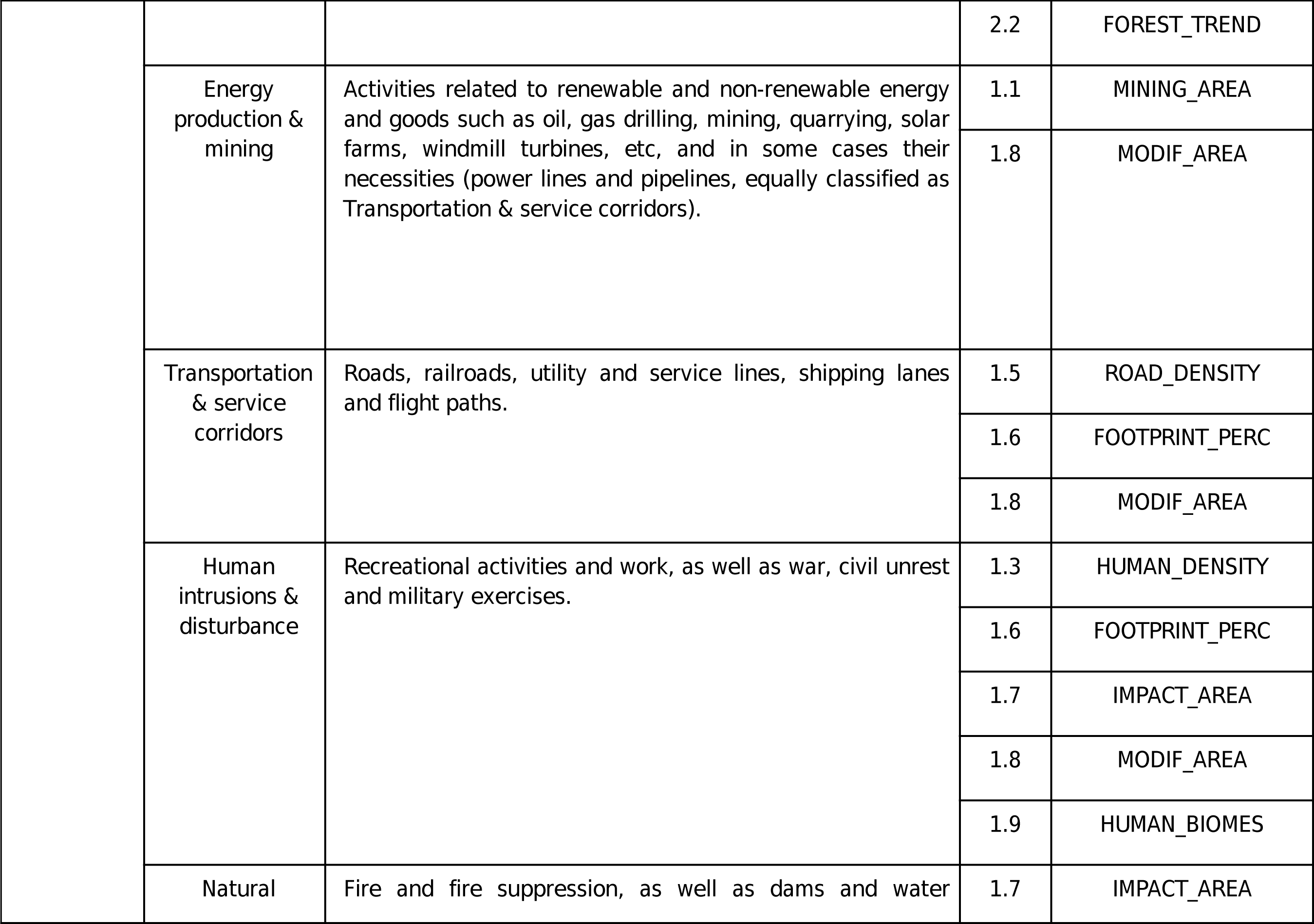
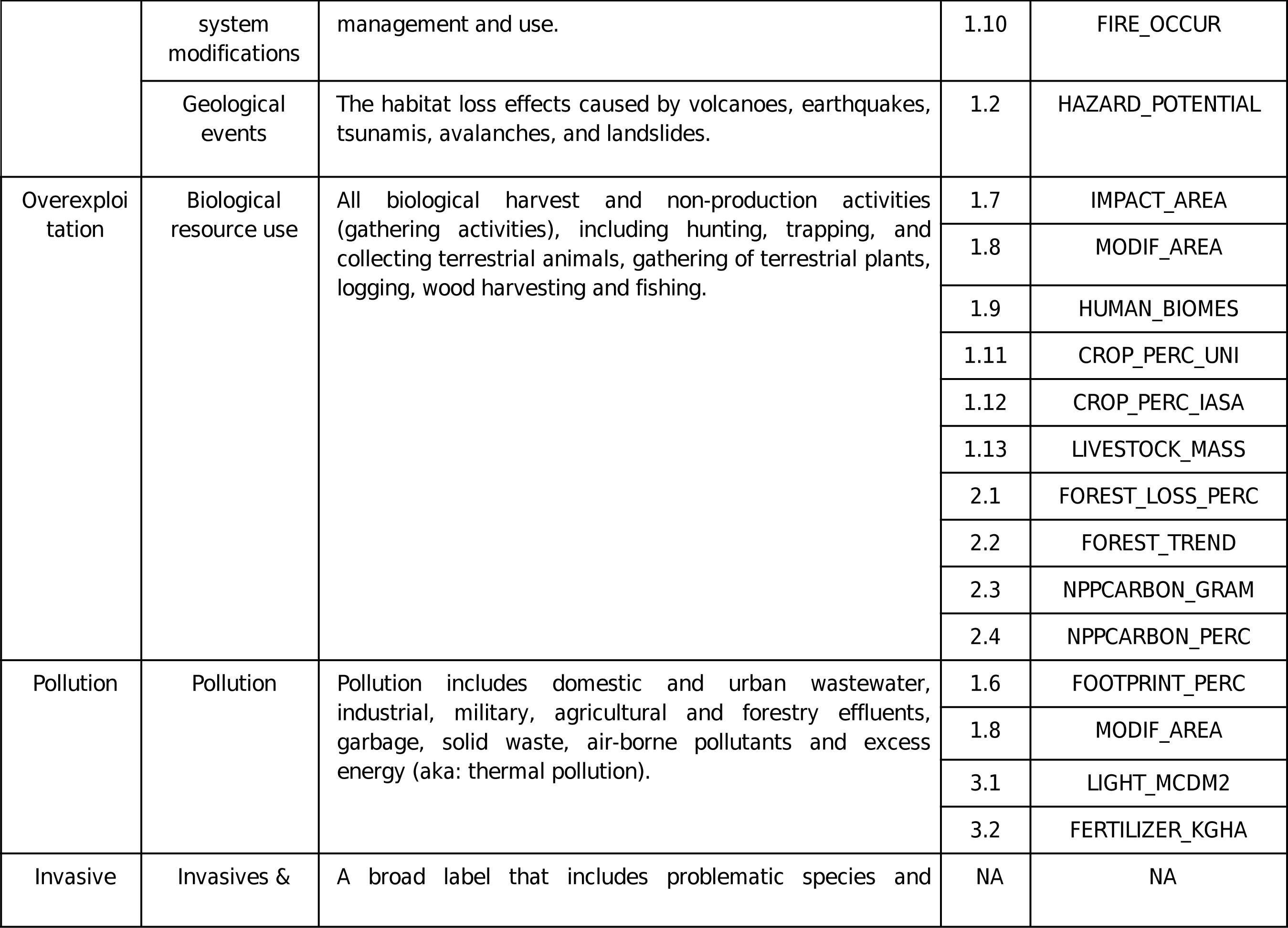
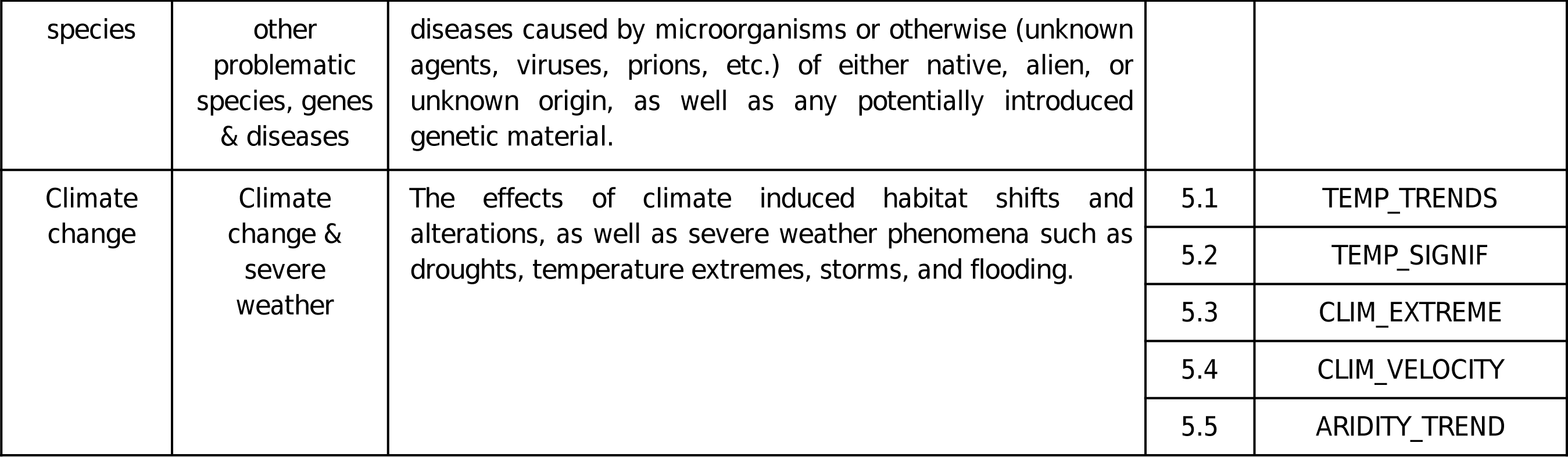
Threat categories associated with each SPECTRE layer currently included in the database (see Table 2), as defined by the Convention on Biological Diversity and the International Union for the Conservation of Nature (IUCN).

All layers cover a time-span between 1980 and 2020 according to the availability of data. Not all variables extend throughout the entire date range. However, we anticipate that threats will exhibit spatial and temporal correlations on a global scale. This means that the regions heavily impacted by a particular threat in one year or decade are likely to have high correlations with the regions most affected in other years or decades by the same threat. Therefore, layers covering different time spans may still be directly comparable.

In this initial phase of the database, we made the decision not to include sets of the same layer covering different periods. This means that for most threats, we do not have data available for different time periods in the database. However, there are other products like human footprint indexes (Keys, Barnes & Carter, 2021; Mu *et al*., 2022) and Global Forest Watch (2014) that do provide limited data for certain variables across different time periods. It is important to note that although this data is not currently available in SPECTRE, we do have plans to include it in future releases of the database. The goal is to expand the availability of data for different periods and to provide a more comprehensive resource for users.

The criteria for inclusion of layers in SPECTRE included data being: 1) publicly available, with a licence that allowed open sharing under a Creative Commons CC BY 4.0 licence or similar public domain or non-commercial licences; 2) mostly global in its intent and scope (see exceptions under post-processing); 3) with a minimum resolution of 50 x 50 km; and 4) the best available products for each threat category according to our own judgement. We excluded data not directly associated with threats, e.g, land use products with a greater diversity of classes than potential threats like cropland or urban areas.

All layers in SPECTRE were created using R (R Core Team, 2023), namely the package terra (Hijmans, Bivand, Pebesma, & Sumner, 2023) and the software QGIS, version 3.10.11 (QGIS Association, 2021). All layers were encoded as FLT4S, reclassified to have −3.4×10^38^ as their NA value and finally masked areas considered as NA in WorldClim. Layers requiring either resampling or reprojection were transformed using the bilinear method for continuous data and the nearest neighbour method for categorical data (Arif & Akbar, 2005; ArcGIS Pro, 2023).

### 2.2 Data transformation

Each layer was transformed to the same resolution, datum and projection as WorldClim (Fick & Hijmans, 2017) so that both databases can be used simultaneously without further adjustments from the user. This implied different manipulation of data to convert every layer to a 30 sec resolution (circa 1 km^2^ at the equator) covering the extent between longitudes - 180 and 180 and latitudes −60 and 90. The temporal scope varied between 1980 and 2020 depending on the availability of information, although some layers (i.e: *HAZARD_POTENTIAL*) use older historical information. We also provide metadata for all layers, including the original citations (Table 2). Across all layers, larger values represent larger impact, with negative values representing a possibly positive impact (e.g. negative forest loss represents a gain in forest area).

**TABLE 2.**
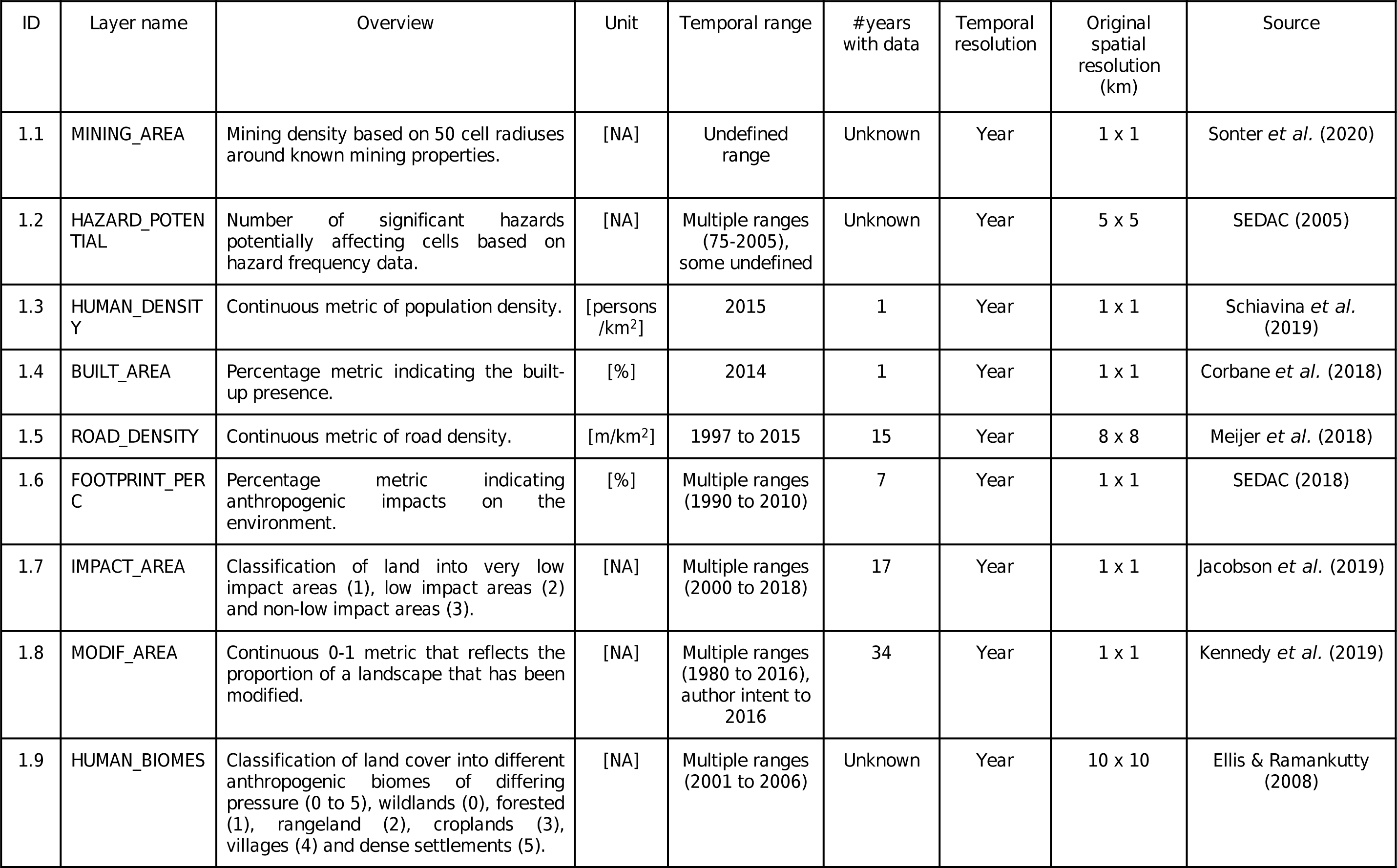
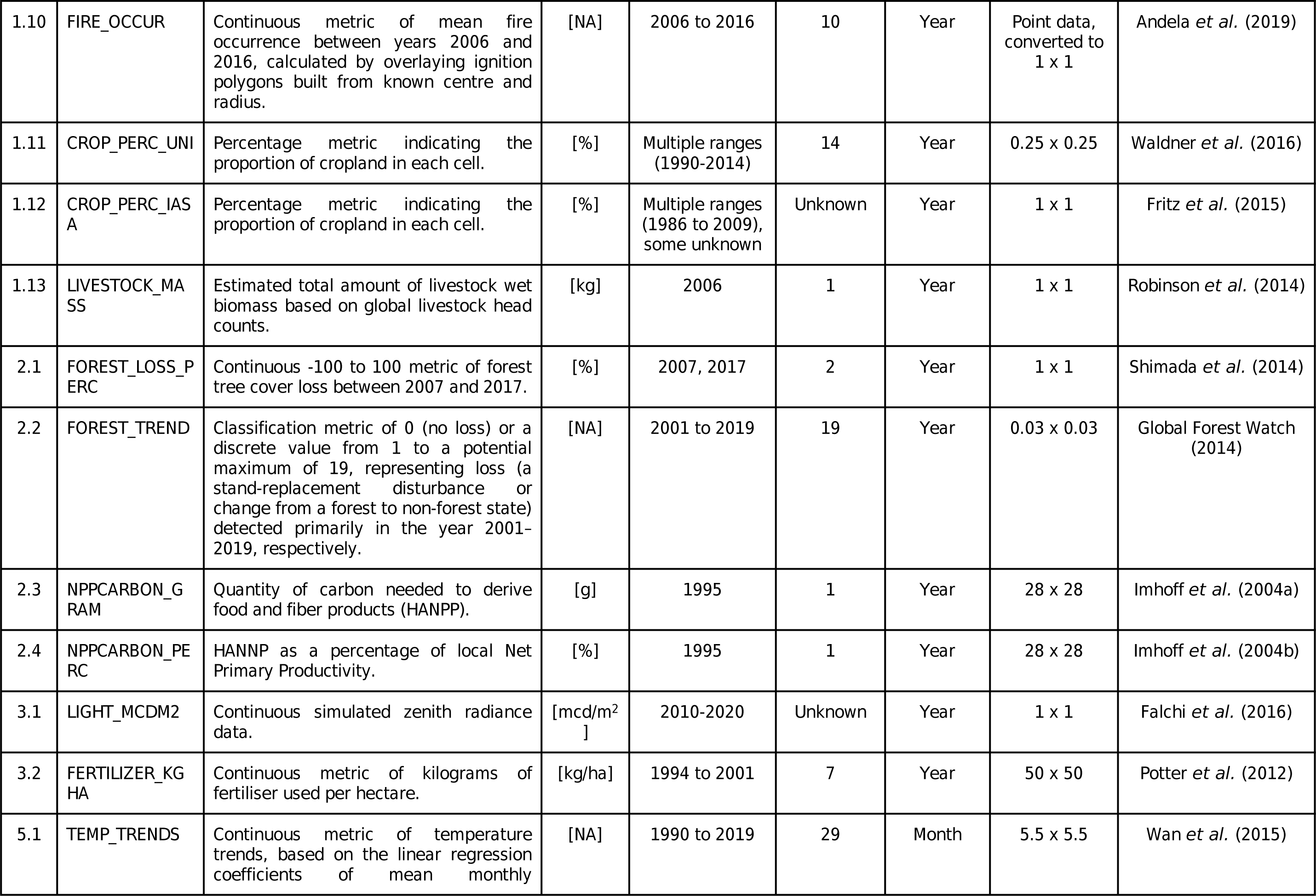
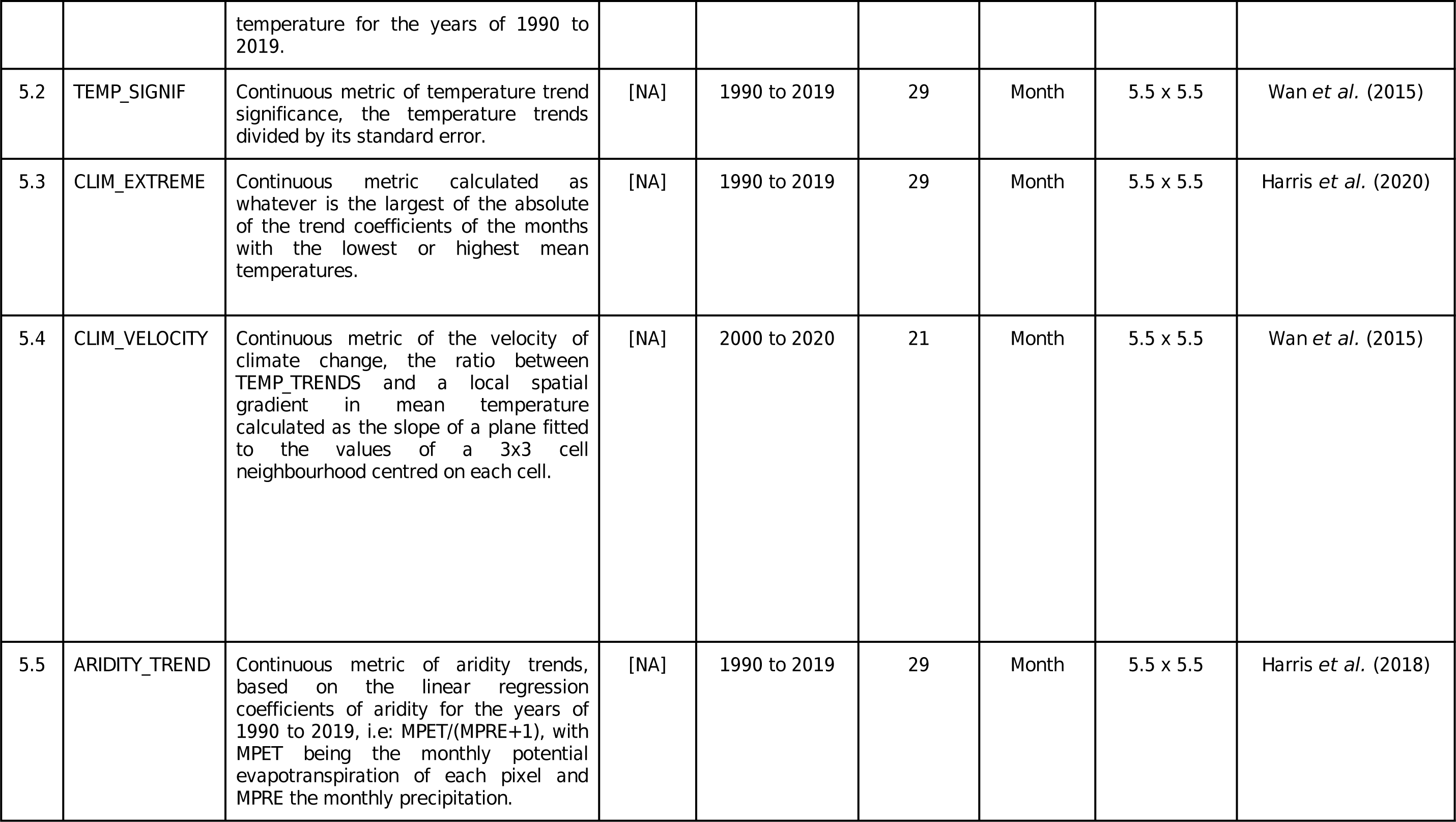
SPECTRE GIS original layer details.

We took measures to reduce incongruence between raster layers, due to missing or out of bounds data associated with water bodies. For this we used the Level 1 vector layer of the Global Lakes and Wetlands Database (Lehner & Döll, 2004), which contains the shoreline polygons of the world’s largest lakes and reservoirs. We then selected all bodies of water with an area larger than 10000 km^2^, rasterized the file and masked our WorldClim template with it, assigning all corresponding pixels as NA.

#### 2.2.1 General procedures

A vast proportion of layers needed only few processing steps, such as resampling to the chosen standard (*FOREST_TREND, FOOTPRINT_PERC*, *FERTILIZER_KGHA*, *LIGHT_MCDM2*, *ROAD_DENSITY, CROP_PERC_UNI*, *CROP_PERC_IIASA*, *HUMAN_BIOMES*, *NPPCARBON_GRAM*, *NPPCARBON_PERC),* reprojection (*HUMAN_DENSITY*, *BUILT_AREA*, *MODIF_AREA*) or masking to terrestrial areas (*HAZARD_POTENTIAL*).

*FOREST_TREND* was created with forest loss data (Global Forest Watch 2014), available for download in segments. These were collected and individually rescaled. *FOOTPRINT_PERC* (SEDAC, 2018) is an index percentage metric indicating anthropogenic impacts on the environment in 2009, created by collating data on human population density, human land use, built-up areas, nighttime lights, land use/land cover and human access (coastlines, roads, railroads, navigable rivers) with a discriminate scoring system. *FERTILIZER_KGHA* (Potter, Ramankutty, Bennett & Donner, 2012) is a continuous metric of nitrogen fertiliser nutrients applied to croplands (with variation in the number of different nutrients, depending on the information the authors possessed) in kilograms of fertiliser used per hectare and collates both fertiliser data as well as geospatial harvested area data between 1994 and 2001. *LIGHT_MCDM2* (Falchi *et al*., 2016) is a measure of continuous simulated zenith radiance data, taking into account all sources contributing to it in a radius of 200 km (representing the impact of medium to long range effects). *ROAD_DENSITY* (Meijer, Huijbregts, Schotten. & Schipper, 2018) is a continuous metric of road density in metres per square kilometre, resulting from the collation of 60 geospatial datasets on road infrastructure with more global coverage than similar products. *CROP_PERC_UNI* and *CROP_PERC_IIASA* (Fritz *et al*., 2015; Waldner *et al*., 2016) are both percentage metrics indicating the proportion of cropland in each cell, differing in temporal grain (1990 to 2014 with an author intent towards a 2005 baseline year) and methodology (adequacy analysis and crowd-sourcing & agreeableness, respectively). *HUMAN_BIOMES* (Ellis & Ramankutty, 2008) is a land cover layer, classifying terrain into different anthropogenic biomes of increasing anthropogenic pressure. The original dataset (see table S1) classified cells into 20 different categories which we reclassified into 6: wildlands (0), forested (1), rangeland (2), croplands (3), villages (4) and dense settlements (5).

*NPPCARBON_GRAM* and *NPPCARBON_PERC* (Imhoff *et al*., 2004a; Imhoff *et al*., 2004b) are continuous metrics of the Human Appropriation of Net Primary Productivity (HANPP) in 1995, expressed respectively as the quantity of carbon needed to derive food (vegetal foods, meat, milk, eggs) and fibre products and the percentage of local Net Primary Productivity appropriated. In the case of *NPPCARBON_PERC*, it is not uncommon for cells to frequently have values upward of 100% as urbanised areas use resources beyond their potential primary production.

*HUMAN_DENSITY* (Schiavina *et al*., 2019) and *BUILT_AREA* (Corbane *et al*., 2018) are continuous metrics of human density for the year 2015, expressed in people per km^2^ and percentage of built-up presence, respectively. Lastly, *MODIF_AREA* (Kennedy, Oakleaf, Theobald, Baruch-Mordo & Kiesecker, 2019) is a continuous metric between 0 and 1 that reflects the proportion of a landscape that has been modified by collating remote sensing imagery and ground inventory data from 1980 to 2016 (with a median year of 2016) on a group of 5 anthropogenic stressors: 1) human settlement (population density & built-up areas), 2) agriculture (cropland & livestock), 3) transportation (major roads, minor roads, two tracks (a kind of two direction dirt road created through pedestrian and vehicle passage) & railroads), 4) mining and energy production (mining, oil wells & wind turbines) and 5) electrical infrastructure (powerlines, nighttime lights).

Additionally, two layers also needed to be masked to terrestrial areas: *HAZARD_POTENTIAL* and *MINING_AREA*. *HAZARD_POTENTIAL* (SEDAC, 2005) is a discrete metric of the number of significant hazards (a.k.a: earthquakes, volcanoes, landslides, floods, droughts & cyclones) potentially affecting cells based on hazard frequency data, based on threat analyses of high temporal grain. *MINING_AREA* is a discrete metric of the number of mines (pre-operational, operational, closed) in a 50 cell radius, considering both exploration of materials critical to renewable energy technology and infrastructure and other materials.

#### 2.2.2 Further transformations

Some of the layers present in SPECTRE required extensive processing of the source data: *IMPACT_AREA*, *FOREST_LOSS_PERC, LIVESTOCK_MASS*, *FIRE_OCCUR*, *TEMP_TRENDS*, *TEMP_SIGNIF*, *CLIM_EXTREME*, *CLIM_VELOCITY* and *ARIDITY_TREND*.

In *IMPACT_AREA* (Jacobson, Riggio, Tait & Baillie, 2019), the original data was composed of two binary rasters which were combined in a single layer representing very low (1), low (2) and high (3) impact. The final result was reprojected to our standard. Similarly, *FOREST_LOSS_PERC* (Shimada *et al*., 2014) is the result of two rasters of tree cover proportion (0-1), for the years 2007 and 2017, the latter being subtracted from the former. The end result is a raster with positive percentages being tree cover losses and negative percentages being tree cover gains.

*LIVESTOCK_MASS* was created using a similar methodology to that described in Bar-On, Phillips & Milo (2018). The average wet biomass values present in Dong, Mangino, Mcallister & Have (2006) were used to infer average wet biomass values for developed, transitioning and developing economies and the inferred values were combined with a country polygon vector map (Natural Earth, 2018) and a classification of world economies (United Nations, 2006) and then applied to 2006 livestock count rasters from the Gridded Livestock of the World (GLW2, Robinson *et al*., 2014) for cattle, chickens, ducks, goats, pigs and sheep, to estimate total livestock biomass.

*FIRE_OCCUR* is a fire occurrence layer created by taking point ignition data (Andela *et al*., 2019) for the years of 2006 to 2016, approximately 1 million points per year, and for each point drawing a polygon using the ignition’s radius.

Five layers for climate change: *TEMP_TRENDS*, *TEMP_SIGNIF*, *CLIM_EXTREME*, *CLIM_VELOCITY* and *ARIDITY_TREND*, were constructed based on methodology described in Bowler *et al*. (2020), using two datasets, Wan, Hook & Hulley (2015) and Harris, Jones & Osborn (2020). All layers used monthly data collected between 1990 and 2019. *TEMP_TRENDS* is a continuous metric of temperature trends, based on the linear regression coefficients of mean monthly temperature. Similarly, *TEMP_SIGNIF* is a continuous metric of temperature trend significance, the temperature trends divided by its standard error. *CLIM_EXTREME* is a continuous metric calculated as whatever is the largest of the absolutes of the coefficients of two linear regressions, one of the month with the lowest mean temperatures, another of the month with the highest mean temperatures.

*CLIM_VELOCITY* is a continuous 0-1 metric of the velocity of climate change calculated as the ratio between two normalised features, *TEMP_TRENDS* and a local spatial gradient in mean temperature. This spatial gradient uses MODIS/Terra Land Surface Temperature (Wan *et al*., 2015) and is calculated through the average maximum technique, defined as the slope of a plane fitted to the values of a 3×3 cell neighbourhood centred on a pixel (ArcGIS Resources, 2021). This reflects the speed at which an organism would have to move to track its current climatic range. To our knowledge two studies have calculated climate velocity at a global scale. Loarie *et al*. 2009 used the WorldClim Version 1.4 Annual Mean Temperature and Total Annual Precipitation bioclimatic variables to compute climate velocity for 2050 to 2100, based on the emissions scenario temporal gradient present in the Special Report on Emissions Scenarios (SRES) A1B. It has several differences to our layer, which is expected given both the different temporal range as well as the diluted effects of temperature trends over three estimates (upper, mean, lower), accentuating patterns characteristic of the local spatial gradient used. More recently, Asamoah *et al*. (2022) have also produced climate velocity layers for both historic data and projected climate scenarios. However, given both the scarce code distributed in the original publication and the regional grassroots nature of the climate data used (CORDEX), we were unable to replicate what was calculated. Despite that, we note patterns suggesting a similar local spatial gradient.

The fifth layer, *ARIDITY_TREND*, is a continuous metric based on the linear regression coefficients of aridity, which is given by the expression:

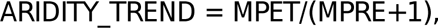

with MPET being the monthly potential evapotranspiration of each pixel and MPRE the monthly precipitation.

## 3 DATA ACCESSIBILITY & USAGE NOTES

SPECTRE is publicly accessible through several services. The Finnish IT Centre for Scientific Computing (CSC 2023) spatial data download service, Paituli (https://paituli.csc.fi/) allows for easy download of all layers in a shared zipped folder. All data can also be accessed through the *gecko* R package (Branco, Correia & Cardoso, 2023). *gecko* is a collection of geographical analysis functions aimed primarily at ecology and conservation science studies, focusing on processing of both point and raster data. It also allows WMS and WCS request capabilities, making it a seamless transition from SPECTRE data access to processing. Lastly, a dashboard allowing quick visualisation and download is also available at the project page (https://biodiversityresearch.org/spectre/).

SPECTRE inherits current data availability and research biases, with entries being overrepresentative of those threats potentially categorised under habitat loss and fragmentation and overexploitation. Additionally, SPECTRE currently has layers representing threats fitting all categories of the classification system proposed (Table 1) except for “*Invasives & other problematic species, genes & diseases*”. To the extent of our knowledge there is no geospatial information representing the impact of invasive or alien species on local ecosystems that meets our criteria for inclusion in either range or spatial resolution. Furthermore, we are also missing data on some aspects of categories considered. For example, under the IUCN threat “*Natural system modifications*’’ we have data on fire occurrence which fits the current definition of “*Fire and fire suppression* [..]” but we do not have data on dams or water management and its use, also aspects of this category.

The layers also have specific limitations, namely the error incurred through upscaling of the source data and the temporal and spatial grain of the sources used, limiting potential usage of the end product. It is however, to the full extent of our knowledge, the best available information. On some occasions, layers have missing areas, often coastal and insular regions or even entire large geographical areas (often Greenland and other arctic locations, Fig. 2). Several options for the interpolation of these points are being weighed in and may be released in a future update alongside the current version with no interpolation.

**FIGURE 2.**
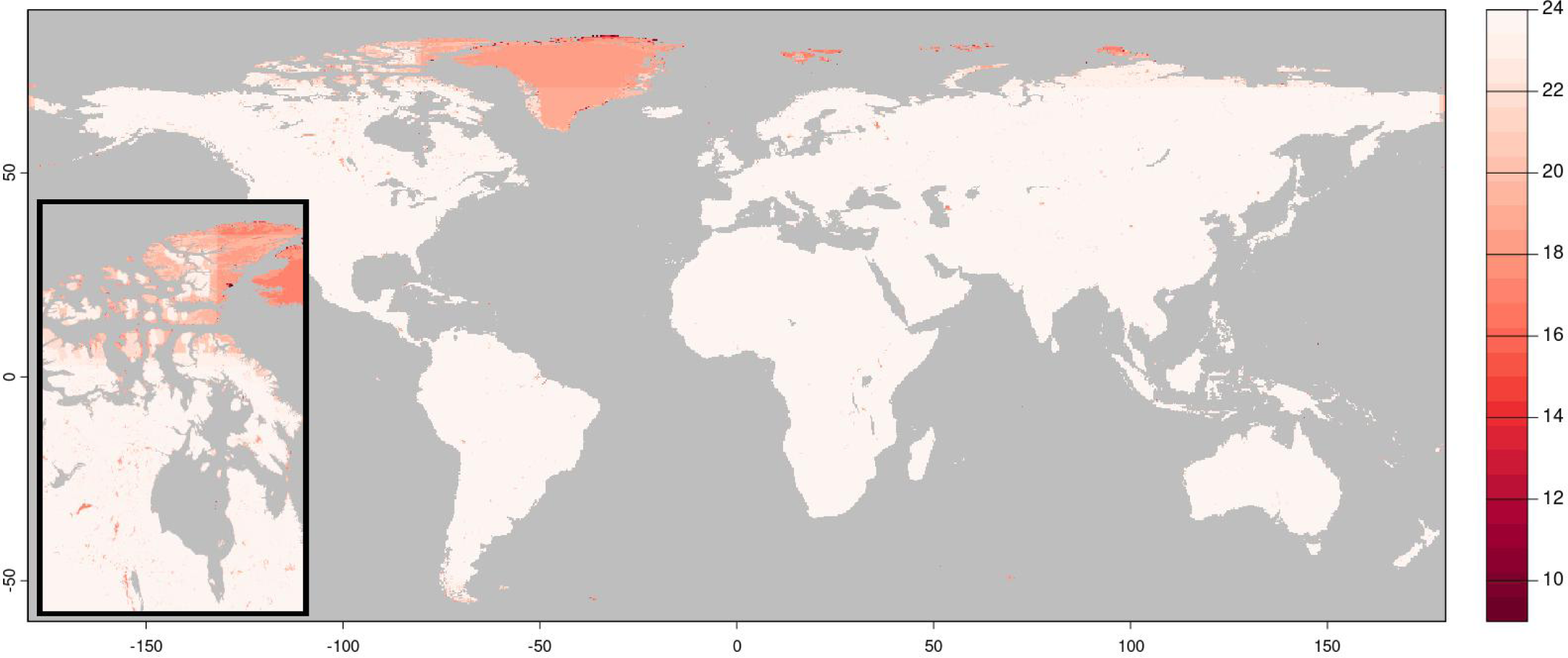
Global map representing the number of layers with information on a given pixel in SPECTRE. On the bottom left corner a close up of the Hudson bay and surrounding areas, highlighting the greater lack of data in higher latitudes and coastal regions.

## ACKNOWLEDGMENTS

V.V.B. and P.C were supported by the Kone Foundation, Finland. L.C. was supported by FCT, Portugal through LASIGE Research Unit, refs. UIDB/00408/2020 and UIDP/00408/2020. We would like to thank Victor Cazalis and Luca Santini for their feedback on the SPECTRE dashboard.

## Notes

https://biodiversityresearch.org/spectre/

### Competing Interest Statement

The authors have declared no competing interest.

### Summary of Updates

author affiliations updated

## REFERENCES

Andela, N., Morton, D. C., Giglio, L., Paugam, R., Chen, Y., Hantson, S.,…Randerson, J. T. (2019). The global fire atlas of individual fire size, duration, speed and direction. Earth System Science Data, 11(2), 529–552. DOI: 10.5194/essd-11-529-2019

ArcGIS Pro (2023). Resample function. Esri. https://pro.arcgis.com/en/pro-app/3.0/help/analysis/raster-functions/resample-function.htm [Accessed 24 November 2023].

ArcGIS Resources (2021). How Slope works. Esri. https://resources.arcgis.com/en/help/main/10.1/index.html#//009z000000vz000000 [Accessed 24 November 2023].

Arif, F. & Akbar, M. (2005). Resampling air borne sensed data using bilinear interpolation algorithm. IEEE International Conference on Mechatronics, 2005, 62–65. DOI: 10.1109/ICMECH.2005.1529228

Asamoah, E. F., Di Marco, M., Watson, J. E. M., Beaumont, L. J., Venter, O. & Maina, J. M. (2022). Land-use and climate risk assessment for Earth’s remaining wilderness. Current Biology, 32(22), 4890–4899. DOI: 10.1016/j.cub.2022.10.016

Bar-On, Y., Phillips, R. & Milo, R. (2018). The biomass distribution on Earth. Proceedings of the National Academy of Sciences, 115(25), 6506–6511. DOI: 10.1073/pnas.1711842115

Bowler, D., Bjorkman, A., Dornelas, M., Myers-Smith, I., Navarro, L., Niamir, A.,…Bates, A. (2020). Mapping human pressures on biodiversity across the planet uncovers anthropogenic threat complexes. People and Nature, 2(2), 380–394. DOI: 10.1002/pan3.10071

Branco, V. V., Correia, L. & Cardoso, P. (2023). gecko: Geographical Ecology and Conservation Knowledge Online. R package version 1.0.0. https://cran.r-project.org/package=gecko

Convention on Biological Diversity (2020a). Aichi Biodiversity Targets. CBD. https://www.cbd.int/sp/targets/ [Accessed 12 November 2020].

Convention on Biological Diversity (2020b). Global Biodiversity Outlook 5. CBD. https://www.cbd.int/gbo/gbo5/publication/gbo-5-en.pdf [Accessed 12 November 2020].

Convention on Biological Diversity (2023). Kunming-Montreal Global Biodiversity Framework. CBD. https://www.cbd.int/gbf/ [Accessed 15 June 2023].

Corbane, C., Florczyk, A., Pesaresi, M., Politis, P., & Syrris, V. (2018). GHS built-up grid, derived from Landsat, multitemporal (1975-1990-2000-2014)*. European Commission*, Joint Research Centre (JRC). DOI: 10.2905/jrc-ghsl-10007

CSC (2023). IT Center for Science. Available at: https://www.csc.fi/ [Accessed 13 June 2023].

Dong, H., Mangino, J., Mcallister, T. & Have, D. (2006). Emissions from livestock and manure management (Intergovernmental Panel on Climate Change) https://www.ipcc-nggip.iges.or.jp/public/2019rf/vol4.html

Dornelas, M., Antão, L. H., Moyes, F., Bates, A. E., Magurran, A. E., Adam, D.,…Zettler, M. L. (2018). BioTIME: A database of biodiversity time series for the Anthropocene. Global Ecology And Biogeography, 27(7), 760–786. DOI: 10.1111/geb.12729

Ellis, E.C. & Ramankutty, N. (2008) Putting people in the map: anthropogenic biomes of the world. Frontiers in Ecology and the Environment, 6(8): 439–447. DOI: 10.1890/070062

Falchi, F., Cinzano, P., Duriscoe, D., Kyba, C. C. M., Elvidge, C. D., Baugh, K.,…Furgoni, R. (2016). The new world atlas of artificial night sky brightness. Science Advances, 2(6). DOI: 10.1126/sciadv.1600377

Fick, S. E., & Hijmans, R. J. (2017). WorldClim 2: new 1-km spatial resolution climate surfaces for global land areas. International Journal of Climatology, 37(12), 4302–4315. DOI: 10.1002/joc.5086

Fritz, S., See, L., McCallum, I., You, L., Bun, A., Moltchanova, E.,…Obersteiner, M. (2015). Mapping global cropland and field size. Global Change Biology, 21(5), 1980–1992. DOI: 10.1111/gcb.12838

Global Biodiversity Information Facility (2022). Global Biodiversity Information Facility. GBIF. https://www.gbif.org/ [Accessed 26 May 2022].

Global Forest Watch (2014). World Resources Institute. http://www.globalforestwatch.org/ [Accessed 24 November 2023].

Harris, I. C., Jones, P. D. & Osborn, T. (2020). CRU TS4.04: Climatic Research Unit (CRU) Time-Series (TS) version 4.04 of high-resolution gridded data of month-by-month variation in climate (Jan. 1901 - Dec. 2019). Centre for Environmental Data Analysis, https://catalogue.ceda.ac.uk/uuid/89e1e34ec3554dc98594a5732622bce9

Hijmans, R. J., Bivand, E., Pebesma, E. & Sumner, M. D. (2023). terra: Spatial Data Analysis. R package version 1.7-29. https://CRAN.R-project.org/package=terra

Hudson, L. N., Newbold, T., Contu, S., Hill, S. L. L., Lysenko, I., De Palma, A.,…Purvis, A. (2016) ‘The database of the PREDICTS (Projecting Responses of Ecological Diversity in Changing Terrestrial Systems) project’. Ecology and Evolution, 7(1), 145–188. DOI: 10.1002/ece3.2579.

Imhoff, M. L., Bounoua, L., Ricketts, T., Loucks, C., Harriss, R. & Lawrence, W. T. (2004a). Global Patterns of HANPP, v1 (1995). NASA Socioeconomic Data and Applications Center. DOI: 10.7927/H44Q7RWV

Imhoff, M. L., Bounoua, L., Ricketts, T., Loucks, C., Harriss, R. & Lawrence, W. T. (2004b). HANPP as a Percentage of Net Primary Productivity, v1 (1995). NASA Socioeconomic Data and Applications Center. DOI: 10.7927/H44Q7RWV

IUCN (2023). Threats classification scheme (version 3.3). IUCN Redlist. https://www.iucnredlist.org/about/barometer-of-life

Jacobson, A. P., Riggio, J., Tait, A. M. & Baillie, J. E. M. (2019), Global areas of low human impact (‘Low Impact Areas’) and fragmentation of the natural world. Dryad. DOI: 10.5061/dryad.z612jm67g

Kennedy, C. M., Oakleaf, J. R., Theobald, D. M., Baruch-Mordo, S., & Kiesecker, J. (2019). Managing the middle: A shift in conservation priorities based on the global human modification gradient. Global Change Biology, 25(3), 811–826. DOI: 10.1111/gcb.14549

Keys, P.W., Barnes, E.A. & Carter, N.H. (2021) ‘A machine-learning approach to human footprint index estimation with applications to sustainable development’, Environmental Research Letters, 16(4), p. 044061. DOI: 10.1088/1748-9326/abe00a.

Lehner, B., & Döll, P. (2004). Development and validation of a global database of lakes, reservoirs and wetlands. Journal of Hydrology, 296(1-4), 1–22. DOI: 10.1016/j.jhydrol.2004.03.028

Loarie, S. R., Duffy, P. B., Hamilton, H., Asner, G. P., Field, C. B. & Ackerly, D. D. (2009). The velocity of climate change. Nature, 462, 1052–1055. DOI: 10.1038/nature08649

Meijer, J. R., Huijbregts, M. A. J., Schotten, K. C. G. J. & Schipper, A. M. (2018). Global patterns of current and future road infrastructure. Environmental Research Letters, 13(6), p.064006. DOI: 10.1088/1748-9326/aabd42

Mu, H., Li, X., Wen, Y., Huang, J., Du, P., Su, W.,…Geng, M. (2022). A global record of annual terrestrial Human Footprint dataset from 2000 to 2018. Sci Data, 9, 176. DOI: 10.1038/s41597-022-01284-8

Natural Earth (2018). Natural Earth Data (vector data v. 4.1.1). https://www.naturalearthdata.com/ [Accessed 24 November 2023].

Potter, P., Ramankutty, N., Bennett., E. M. & Donner, S. D. (2012). Global Fertilizer and Manure, Version 1: Nitrogen Fertilizer Application. NASA Socioeconomic Data and Applications Center. DOI: 10.7927/H4Q81B0R

QGIS Association (2021). QGIS Geographic Information System. QGIS. http://www.qgis.org

R Core Team (2023). R: A Language and Environment for Statistical Computing. R Foundation for Statistical Computing. https://www.R-project.org/

Robinson, T. P., Wint, G. R. W., Conchedda, G., Van Boeckel, T. P., Ercoli, V., Palamara, E.,…Gilbert, M. (2014). Mapping the global distribution of Livestock. PLoS ONE, 9(5). DOI: 10.1371/journal.pone.0096084

Schiavina, M., Freire, S., & MacManus, K. (2019). GHS-pop R2019A - GHS population grid multitemporal (1975-1990-2000-2015). *European Commission*, Joint Research Centre (JRC). DOI: 10.2905/42E8BE89-54FF-464E-BE7B-BF9E64DA5218

SEDAC (2005). Global Multihazard Frequency and Distribution, v1: Natural Disaster Hotspots. NASA Socioeconomic Data and Applications Center. DOI: 10.7927/H45718Z5

SEDAC (2018). Global Human Footprint (Geographic), v3 (1995–2004): Last of the Wild. NASA Socioeconomic Data and Applications Center. DOI: 10.7927/H46T0JQ4

Shimada, M., Itoh, T., Motooka, T., Watanabe, M., Shiraishi, T., Thapa, R., & Lucas, R. (2014). New Global Forest/non-forest maps from Alos Palsar data (2007–2010). Remote Sensing of Environment, 155, 13–31. DOI: 10.1016/j.rse.2014.04.014

Sonter, L.J., Dade, M.C., Watson, J.E.M. & Valenta, R. K. (2020). Renewable energy production will exacerbate mining threats to biodiversity. Nature Communications 11, 4174. DOI: 10.1038/s41467-020-17928-5

United Nations (2006). World Economic Situation and Prospects 2006. United Nations New York. https://www.un.org/en/development/desa/policy/wesp/wesp_archive/2006wesp.pdf [Accessed 16 April 2021].

Waldner, F., Fritz, S., Di Gregorio, A., Plotnikov, D., Bartalev, S., Kussul, N.,…Defourny, P. (2016). A unified cropland layer at 250 m for Global Agriculture Monitoring. Data, 1(1), 3. DOI: 10.3390/data1010003

Wan, Z., Hook, S. & Hulley, G. (2015). MOD11C3 MODIS/Terra Land Surface Temperature/Emissivity Monthly L3 Global 0.05Deg CMG V006. NASA EOSDIS Land Processes DAAC. DOI: 10.5067/MODIS/MOD11C3.006

