## Supplementary material for "SPECTRE: standardized global spatial data on terrestrial SPECies ThREats": Table S1

**Table S1**. Original values and labels present in the dataset by Ellis & Ramankutty (2008) and their corresponding value and labels in *HUMAN_BIOMES.* Given an already established order, we classified according to increasing anthropogenic pressure.

| Anthropogenic biomes of the world | | *HUMAN_BIOMES* | |
|---|---|---|---|
| Value | Label | Value | Label |
| 11 | Urban | 5 | Dense settlements |
| 12 | Dense settlements | | |
| 22 | Irrigated villages | 4 | Villages |
| 23 | Cropped & pastoral villages | | |
| 24 | Pastoral villages | | |
| 25 | Rainfed villages | | |
| 26 | Rainfed mosaic villages | | |
| 31 | Residential irrigated cropland | 3 | Croplands |
| 32 | Residential rainfed mosaic | | |
| 33 | Populated irrigated cropland | | |
| 34 | Populated rainfed cropland | | |
| 35 | Remote croplands | | |
| 41 | Residential rangelands | 2 | Rangelands |
| 42 | Populated rangelands | | |

| | | | |
|---|---|---|---|
| 43 | Remote rangelands | | |
| 51 | Populated forests | 1 | Forested |
| 52 | Remote forests | | |
| 61 | Wild forests | 0 | Wildlands |
| 62 | Sparse trees | | |
| 63 | Barren | | |
